# Altered NPTX2 dynamics associated with impaired cognitive aging

**DOI:** 10.1101/2025.09.15.676156

**Authors:** Rebecca P. Haberman, Ana Delgado, Meifang Xiao, Ming Teng Koh, Ashley A. Becker, Shiyu Ji, Paul F. Worley, Michela Gallagher, Audrey Branch

## Abstract

Changes in synaptic integrity and neural activity homeostasis are hallmarks of brain aging and are closely tied to cognitive outcomes. Yet, defining their relationship across the continuum from normal aging to neurodegenerative disease has proven challenging. Recent research investigating the dynamic changes of neuronal pentraxin 2 (NPTX2, or Narp, neuronal activity-related pentraxin) in cerebrospinal fluid (CSF) of Alzheimer’s disease (AD) subjects supports its promise as a prognostic marker of disease progression, possibly as an expression of synaptic damage related to cognitive impairment. However, studies in human subjects are unable to clearly differentiate age-related and disease-related processes. Here we took advantage of a well-characterized rat model that displays substantial individual differences in hippocampal memory during aging, uncontaminated by slowly progressive, spontaneous neurodegenerative disease. Through this approach, we aimed to interrogate the underlying neural substrates that mediate aging as a uniquely permissive condition and the primary risk for neurodegeneration. We found that successful cognitive aging is associated with an elevation of NPTX2 levels above that found in young or cognitively impaired subjects. Pharmacological engagement of neural activity was sufficient to increase NPTX2 levels in all subjects, but cognitively-impaired aged subjects failed to recruit NPTX2 in response to a hippocampus-dependent memory task. Together the findings demonstrate that changes in NPTX2 are coupled to differential cognitive outcomes of aging, and that successful neurocognitive aging is associated with adaptive upregulation of NPTX2, not simply the persistence of youthful synaptic dynamics.

**Significance Statement:** Although aging is the most prominent risk factor for Alzheimer Disease (AD), the age-dependent processes that disrupt neurophysiological homeostasis leading to neurodegenerative disease remain poorly defined. Alterations in NPTX2, a synaptic protein biomarker for AD, may elucidate aging processes underlying pathological trajectories. We examined NPTX2 in an aging context and identified circuit specific alterations of NPTX2 that are coupled with distinct memory outcomes in aging in the absence of potential confounds of neurodegenerative disease. Greater NPTX2, associated with successful cognitive aging, may reflect coordinated molecular and circuit-level adaptations that sustain memory-relevant hippocampal activity. Development of targets and interventions that promote neuroadaptive network homeostasis, bending the trajectory of aging away from neurodegeneration, are a potentially valuable alternative to current therapeutic strategies.

## Introduction

Advancing age remains the single greatest risk factor for neurodegenerative conditions, specifically Alzheimer’s disease (AD) and other dementias. The changes unfolding in the aging brain are intimately connected to the processes leading to neurodegenerative disease. However, variance in aging outcomes in humans includes both neurodegenerative dementing disease as well as resilience to progressive cognitive decline (Fig. 1A). Consequently, it remains challenging to determine normal aging versus disease-related cognitive trajectories in human clinical studies (1, 2). Furthermore, the appearance of cognitive symptoms in those ultimately diagnosed with AD is highly heterogenous in terms of age of onset and rates of decline, compounding the difficulty in predicting later outcomes on the basis of clinical presentation alone (2). Development of biomarkers for amyloid-beta (Aβ) and tau protein aggregates have improved patient monitoring and outcome prediction (3, 4), but they still only account a small portion of the variance in cognitive impairment in AD (5). This suggests the existence of additional drivers of AD dementia that are not captured by biomarkers of primary AD pathologies including amyloid and tau. Identifying specific early indicators of age-dependent processes that seed neurogenerative diseases provides a critical foundation to classify at risk individuals and develop early interventions.

**Figure 1.**
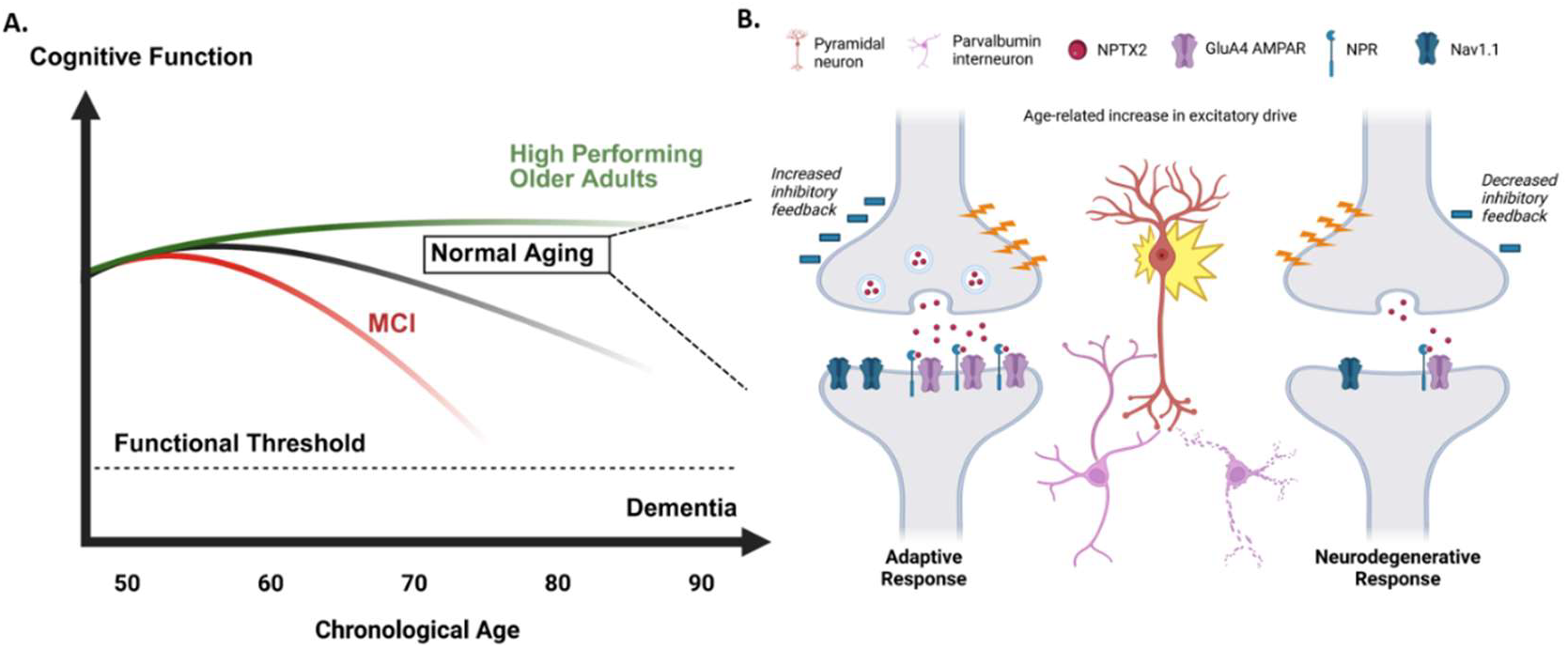
Schematic of NPTX2 in aging and homeostatic control of activity. A) Over the course of chronological aging, cognitive function has a tendency to decline (normal aging, gray line), although some individuals maintain high cognitive function (green line). Subjective memory decline that is worse than expected for one’s age may indicated the beginning of an pathological trajectory, such as prodromal Alzheimer’s disease (red line). B) Aging is associated with alterations in brain activity patterns in key regions mediating memory processes such that neural engagement is accompanied by increased excitatory drive. If left unchecked, this increased excitatory drive results in hyperactivity in neurons engaged in cognitive processing, which ultimately drives pathophysiological changes. Parvalbumin (PV) interneurons are well positioned to maintain homeostatic activity balance in aging neural circuits. In healthy individuals (left), the NPTX2 protein (red) arranges GluA4-containing AMPA receptors (purple) in clusters at these connections via interactions with its receptor NPR (blue). Excitation of presynaptic excitatory neurons strongly activate PV interneurons, resulting in stabilization of Nav1.1 (blue) sodium channels in the PV interneurons which generate electrical signals provide homeostatic negative feedback of other neurons in the circuit. In the brains of individuals on a neurodegenerative trajectory (right), the levels of NPTX2, GluA4 and Nav1.1 are all lower than in healthy individuals, resulting in less inhibitory interneuron activity. Other neurons in the circuit thus become hyperactive, resulting in defects in circuit function and cognitive impairments. We hypothesize that this reduction of NPTX2 is a feature of aging which promotes risk for establishment of early Alzheimer’s disease initiation. Graphic created with BioRender.com.

NPTX2 has shown promise as one such indicator. NPTX2 is a validated biomarker of AD, showing reduced CSF and brain levels in AD patients relative to cognitively normal controls (6) and a tight correlation with cognitive symptoms across both preclinical and clinical disease states up to 15 years prior to onset of cognitive symptoms (7–10). Those clinical studies suggest that reductions of NPTX2 may represent a component of physiological brain aging related to synaptic homeostasis that confers risk for cognitive decline and AD (8, 11). Importantly, NPTX2 is a plasticity-related immediate early gene involved in the regulation of homeostatic negative feedback and stabilization of excitatory synapses onto parvalbumin interneurons (12–15) thus modulating the inhibitory tone of the hippocampus (Fig. 1B). Indeed circuit specific reductions in synaptic integrity and excess neural activity in the medial temporal lobe (MTL) contribute to memory impairment even in the absence of pathological markers of AD (16–20). Thus, NPTX2, as a key synaptic regulator of circuit dynamics, is positioned to modulate processes that are central to the risk for AD conferred by aging itself. At present, available data from studies in humans do not indicate whether NPTX2 reductions depend on neurodegenerative processes or represent a component of aging that may predispose individuals to disease. To determine the role of NPTX2 in the aging brain, without the potential confound of incipient AD, we examined NPTX2 expression in the MTL of aged Long Evans rats, a well-established model of cognitive aging in the absence of neurodegeneration (21–23).

Outbred Long Evans (LE) rats exhibit a naturally occurring variation in cognitive abilities relative to young adults that can be segregated into cohorts based on either impaired or preserved performance on behavioral tasks assessing hippocampal-dependent spatial memory (21). Accordingly, this model allows assessment of cellular and molecular processes underlying cognitive performance in aging apart from neurodegenerative processes that occur in humans. Previous studies have shown that age-dependent circuit alterations associated with impairments in spatial memory in rodents align with those of clinical studies of early amnestic MCI, particularly including altered neuronal excitability in the CA3 and dentate gyrus (DG) subregions of the hippocampus (18–20, 24–26). Furthermore, therapeutic interventions targeting neuronal hyperactivity improve spatial memory performance in rodents (24, 27–29) and also improve memory performance in a hippocampal dependent memory task in clinical studies (30–32).

The present study examines NPTX2 levels in the dorsal hippocampus in the LE rat model of cognitive aging. Results show that NPTX2 levels are elevated in aged rats with intact or unimpaired (AU) spatial memory relative to aged rats with impaired performance (AI). That elevation in NPTX mRNA is also significantly greater than that observed in young adult rats (Y). It is of further interest that AI rats exhibit a normal pharmacological induction of NPTX2 mRNA that corresponds to gene expression markers of neuronal activity levels. In contrast, AI rats do not exhibit an activity-dependent regulation of NPTX2 mRNA in a gene induction protocol using a behavioral memory task. These findings suggest that impaired NPTX2 recruitment in aging may contribute to cognitive decline, resulting in a memory network with reduced resiliency and dynamic range for scaling heterosynaptic homeostasis. Our findings strengthen the assertion that dysregulation of NPTX2 and its associated molecular and cellular processes contribute to impairment due to aging itself as a risk for neurodegeneration and clinical disease.

## Results

### Behavioral Characterization

To determine spatial memory ability, rats were characterized on a hidden platform water maze prior to any manipulations. The primary performance measure, the learning index (LI), is derived from interpolated probe trials throughout the 8-day protocol and has proven a robust measure of spatial learning capability (21). By this measure, low values reflect searching near the platform position and better memory. Aged rats (24 mos) performing within the normative range of young rats are designated aged unimpaired (AU, LI < 240) and those performing worse than young are designated aged impaired (AI, LI > 240), with each group constituting approximately half of the aged cohort. Rats were chosen for each subsequent experiment to represent a range of spatial learning performance and to create balanced groups across treatments (see Fig. S1 for learning index distribution for each experiment).

### Basal NPTX2 expression across MTL

Our initial investigation leveraged a large hippocampal RNA-seq dataset to identify indicators of age and cognitive impairment-related expression differences in NPTX2 and associated genes involved in the neuronal pentraxin homeostatic inhibitory pathway (Fig 2A). Previous work in aged LE rats identified the CA3 subfield of the hippocampus as a region critically affected by age and integral to differing cognitive outcomes (18, 20, 33). Examination of global gene expression profiling in CA3 uncovered correlations between levels of NPTX2, NPTX1 and NPTXR and degree of cognitive impairment in aging (Fig.2A). None of the other genes within the NPTX2 functional network showed a relationship with learning index (LI), although several correlations between genes were observed. An examination of group differences in RNA-seq mRNA reads (Fig. 2B) confirmed a reduction in NPTX2 mRNA levels in AI animals relative to AU subjects (p<0.001, q<0.1). NPTXR and NPTX1 also had decreased expression in AI rats, however this did not meet multiple comparison correction cut-off (q<0.1). GRIA4 (GluR4) mRNA showed no change with cognitive status across aged cognitive groups. Given NPTX2’s status as an immediate early gene, and the most significantly altered among the three neuronal pentraxins in the RNA-seq analysis, we further examined NPTX2 expression in medial temporal lobe (MTL) subfields under homecage conditions as well as treatment conditions that resulted in elevated neuronal activity or hippocampal engagement.

**Figure 2.**
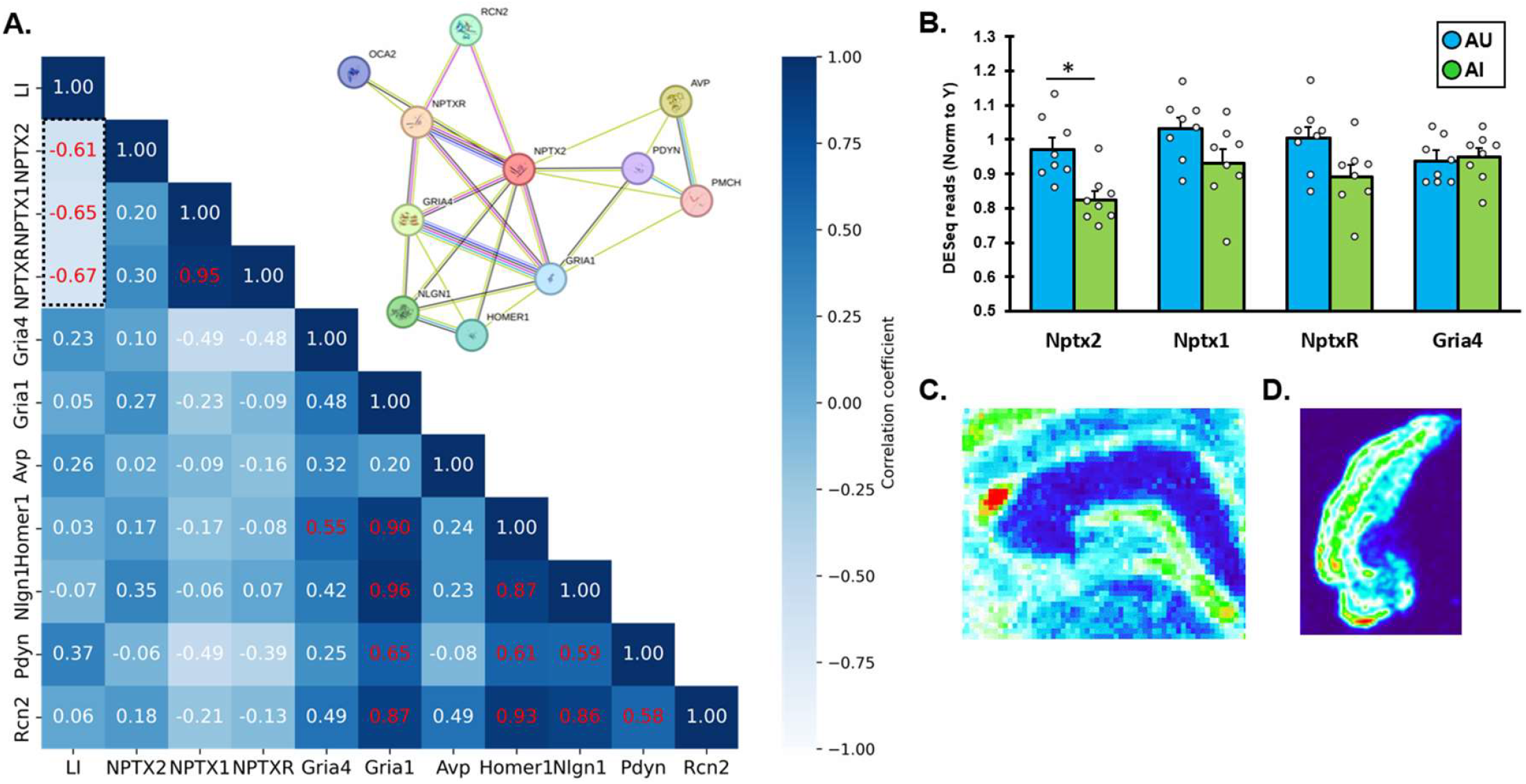
NPTX2 pathway in aged rats. A) The STRING database Protein-Protein Interaction Network (PPIN) for NPTX2; in the inset diagram, nodes indicate known proteins and edges denote protein interactions. DESeq reads for PPIN identified proteins from dissected whole CA3 of aged rats are shown as a correlation matrix, with Pearson r values shown in red font if p-value < 0.05. Reads for NPTX2 and its direct binding partners NPTXR and NPTX1 are found to correlate with learning index. B) DESeq reads for selected proteins in CA3 of aged unimpaired and aged impaired rats (∗ p= 0.01, q < 0.1). C) Representative expression pattern of NPTX2 mRNA in hippocampus and D) entorhinal cortex. Images are pseudocolored micrographs of radiolabeled anti-sense mRNA probe specific for NPTX2 transcripts.

We performed RNA *in situ* hybridization with a series of sections comprising the dorsal hippocampus from Y, AU, and AI rats perfused directly from their homecage. Consistent with the RNA-seq data, we observed reduced basal expression of NPTX2 in AI rats relative to AU in DG, CA3, and CA2 hippocampal subfields but not CA1 (Fig.3A-D; DG: p<0.01, CA3: p<0.000, CA2: p<0.05). This appears to be due to increased expression of NPTX2 in AU rather than reduced expression in AI, as AI levels matched those of Y subjects across all the regions examined (all p>0.12) and AU levels were significantly increased relative to Y (DG: p<0.000, CA2: p<0.01, CA3: p<0.025). Similar to the RNA-seq results (Fig.2), when levels of NPTX2 expression were assessed in aged animals relative to their learning index scores, we observed a significant negative correlation in DG, CA3, and a trend in CA2 (Fig.3A-D) indicating that poorer performance on a MTL dependent task in aged subjects is related to lower levels of NPTX2. The relationship between cognitive status and NPTX2 expression in the MTL was restricted to the hippocampus; investigation of the entorhinal cortices did not reveal an age- or cognition-related alteration in NPTX2 mRNA levels (Fig.3E, F), nor a relationship with cognitive impairment (Fig. 3E, F).

**Figure 3.**
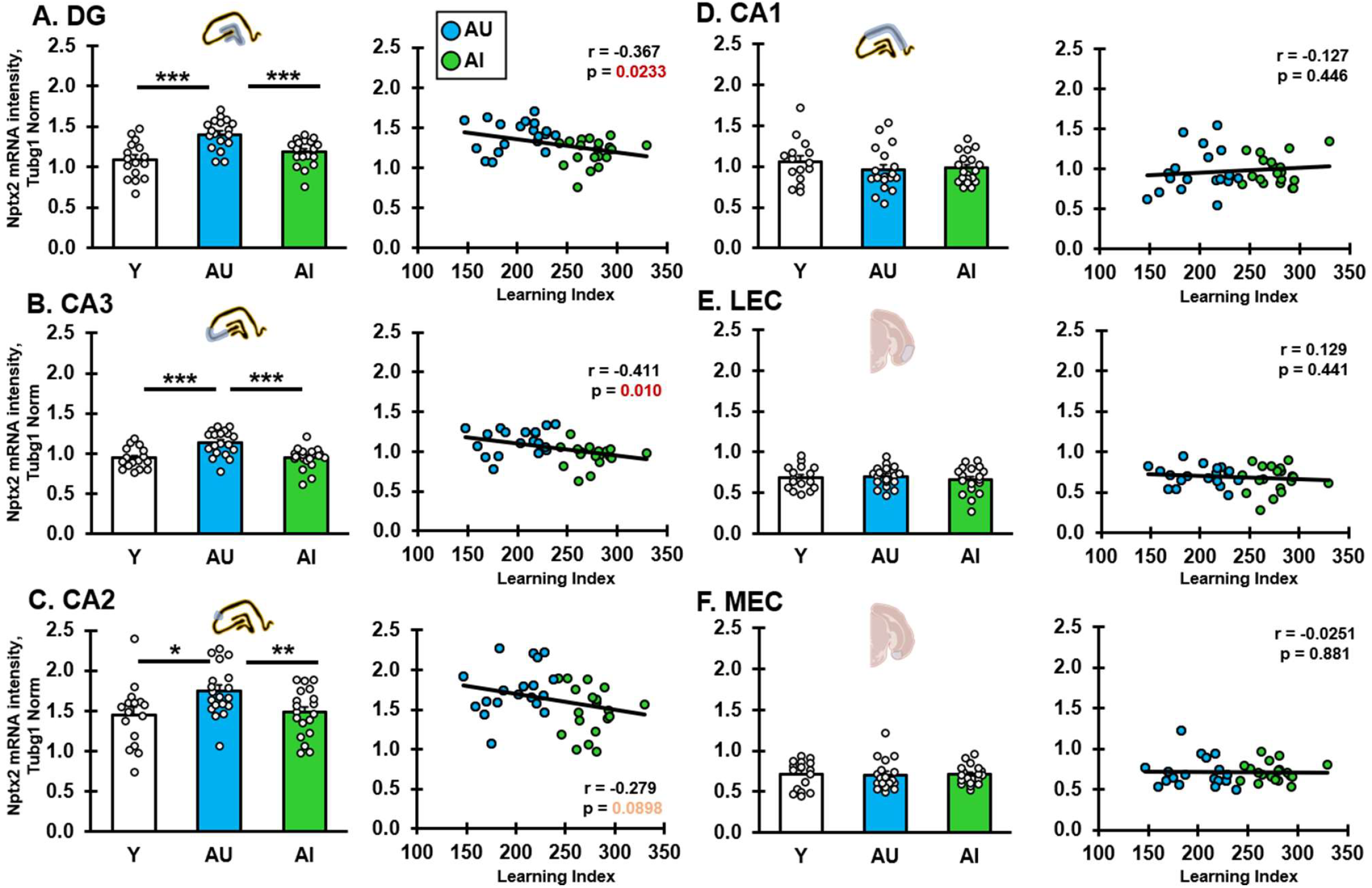
Expression of NPTX2 mRNA in subfields of the medial temporal lobe in young and aged rats. Subfield quantified is indicated by label and inset depicts ROIs used to quantify mRNA signal. All y axis values are NPTX2 mRNA signal intensity normalized to Tubg1 mRNA signal plotted by group or versus learning index for each animal with individual animals shown as data points and bars represent group averages ± SEM. For correlation panels, Pearson r values are indicated in the corner of each graph and all correlations are significant at p < 0.05. Statistical significance for bar plots was determined by t-test as indicted on the graphs: ∗p < 0.05; ∗∗p < 0.01, ∗∗∗p < 0.001. Full statistics are provided in Supp. Fig. 2. Graphic insets created with BioRender.com.

As an immediate early gene, higher levels of NPTX2 are linked to neural activation. In addition, NPTX2 is a positive regulator of both feedforward and feedback inhibition resulting in homeostatic regulation of circuit activity (12, 13). We considered two potential explanations for the elevated NPTX2 expression in AU rats: 1) Elevated NPTX2 mRNA reflects increased inhibitory tone in AU rats specifically in the input areas of the hippocampal circuit, consistent with previous data in this model (26) or 2) Elevated NPTX2 mRNA primes the circuit to rapidly and efficiently encode new information, allowing the AU to retain young-like behavior with a reduced requirement for newly synthesized mRNA. If the elevated NPTX2 contributes to inhibitory tone, we would expect the mRNA to result in increases in protein under the same conditions in which mRNA levels were assessed (i.e. homecage). Western blot analysis of NPTX2 protein in dissected hippocampal subfields identified a negative correlation with learning index (Fig.4A) in the CA1 subfield of the hippocampus, but not in the CA3 or DG subfields (data not shown). Because NPTX2 has been shown to stabilize glutamate receptors at excitatory synapses onto parvalbumin positive interneurons, we examined the levels of GluA4, the primary glutamate receptor subunit involved in this circuit (12, 13, 34). Similar to NPTX2, GluA4 protein showed a negative correlation with learning index in CA1 (Fig.4B). As expected, given the role of NPTX2 in clustering and stabilizing GluA4, we found a positive correlation between NPTX2 and GluA4 protein levels in CA1 across all aged subjects (Fig.4C). Given that CA1 forms the primary output of the hippocampal computational circuit and the high level of NPTX2 mRNA in the upstream CA3 subfield, these correlations support a model by which the NPTX2 produced by CA3 neurons is trafficked to CA1, modulating the ultimate output of the hippocampus. This provides support for a model where elevated baseline levels of NPTX2 mRNA contributes to a greater inhibitory tone in the hippocampal circuit of aged unimpaired subjects.

**Figure 4.**
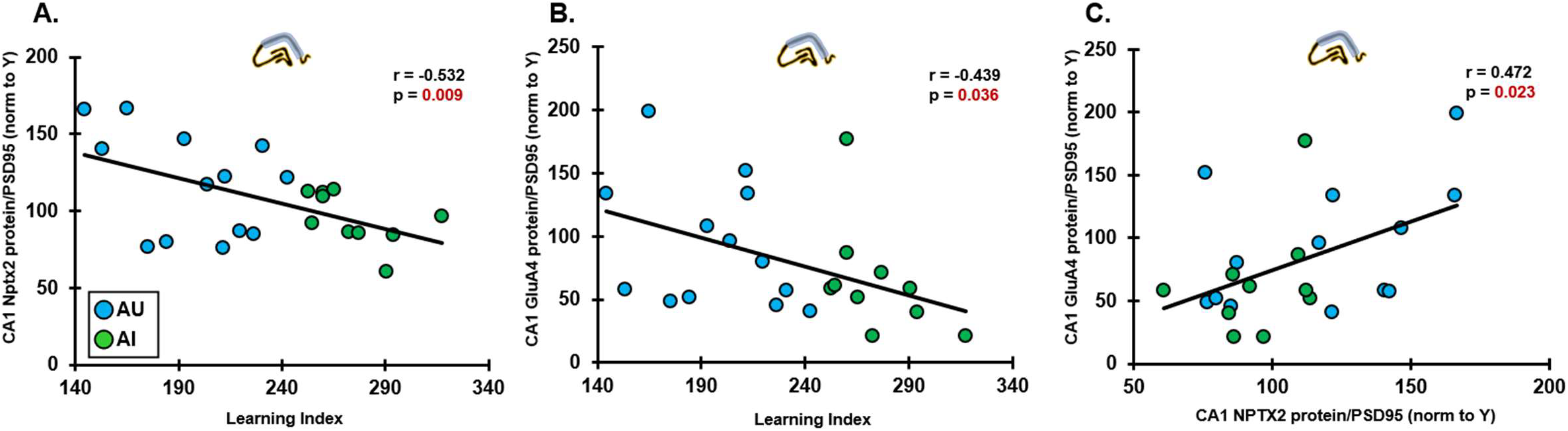
Quantification of NPTX2 and GluA4 protein in CA1. A) NPTX2 protein levels in CA1 were negatively correlated with learning index for aged subjects, indicating that aged animals with worse age-related cognitive impairment have lower levels of NPTX2. B) Levels of GluA4, an AMPA receptor associated enriched on parvalbumin neurons which is stabilized by NPTX2, were also negatively correlated with learning index in aged subjects. C) Levels of NPTX2 and GluA4 proteins levels for individual aged subjects were positively correlated. Data points are values for individual aged animals normalized to young and to PSD95. Pearson r values are indicated in the corner of each graph and all correlations are significant at p < 0.05.

### Activity induced expression

Because the two hypotheses noted above are not mutually exclusive, we further investigated whether elevated NPTX2 acts to prepare neurons for new learning. We examined the ability of aged rats, both AU and AI, to upregulate NTPX2 mRNA in response to 1) broad neuronal activation and 2) behaviorally induced neuronal activation. The ability to respond to neuronal activity is particularly relevant given the documented relationship between NPTX2 expression and neuronal activity (14), and the disruption of E/I balance found in CA3/DG in AI rats (18, 26, 35). To pharmacologically induce global neural activity-dependent gene expression, we administered a subconvulsive dose of pilocarpine and measured DG and CA3 NPTX2 mRNA 1 hour and 4 hours later, a manipulation previously shown to induce immediate early gene expression in young and aged rats (35, 36) (Fig.5A). Both AU and AI rats showed elevated NPTX2 expression at 1 and 4 hours post pilocarpine, consistent with young rats, in all subfields of the hippocampus (Fig. 5B, C; two factor univariate analysis, DG: p<0.005; CA1, CA2, CA3: p<0.000). While treatment with pilocarpine significantly induced NPTX2 expression in all subfields (effect of treatment: DG, CA1, CA2, CA3: p<0.0001), the majority of the induction appeared to occur within the first hour as 1Hr and 4 Hr treatments were both significantly different from homecage (DG, HC vs 1hr p=0.013, HC vs 4hr p<0.001; CA3 HC vs 1Hr and HC vs 4Hr p<0.001) but not different from each other (DG, CA3 1Hr vs 4Hr p>0.25). Furthermore, both DG and CA3 revealed a significant effect of subject group (DG: p=0.033, CA3: p=0.038). The rapid and sustained induction of NPTX2 mRNA in both AU and AI shows that cellular capabilities exist in these animals that allow for activity-dependent regulation of NPTX2 transcription. The timeframe of NPTX2 mRNA induction was consistent with previous studies, which have shown near maximal levels at 1-hour post induction that remain elevated for up to 12 hours (14).

**Figure 5.**
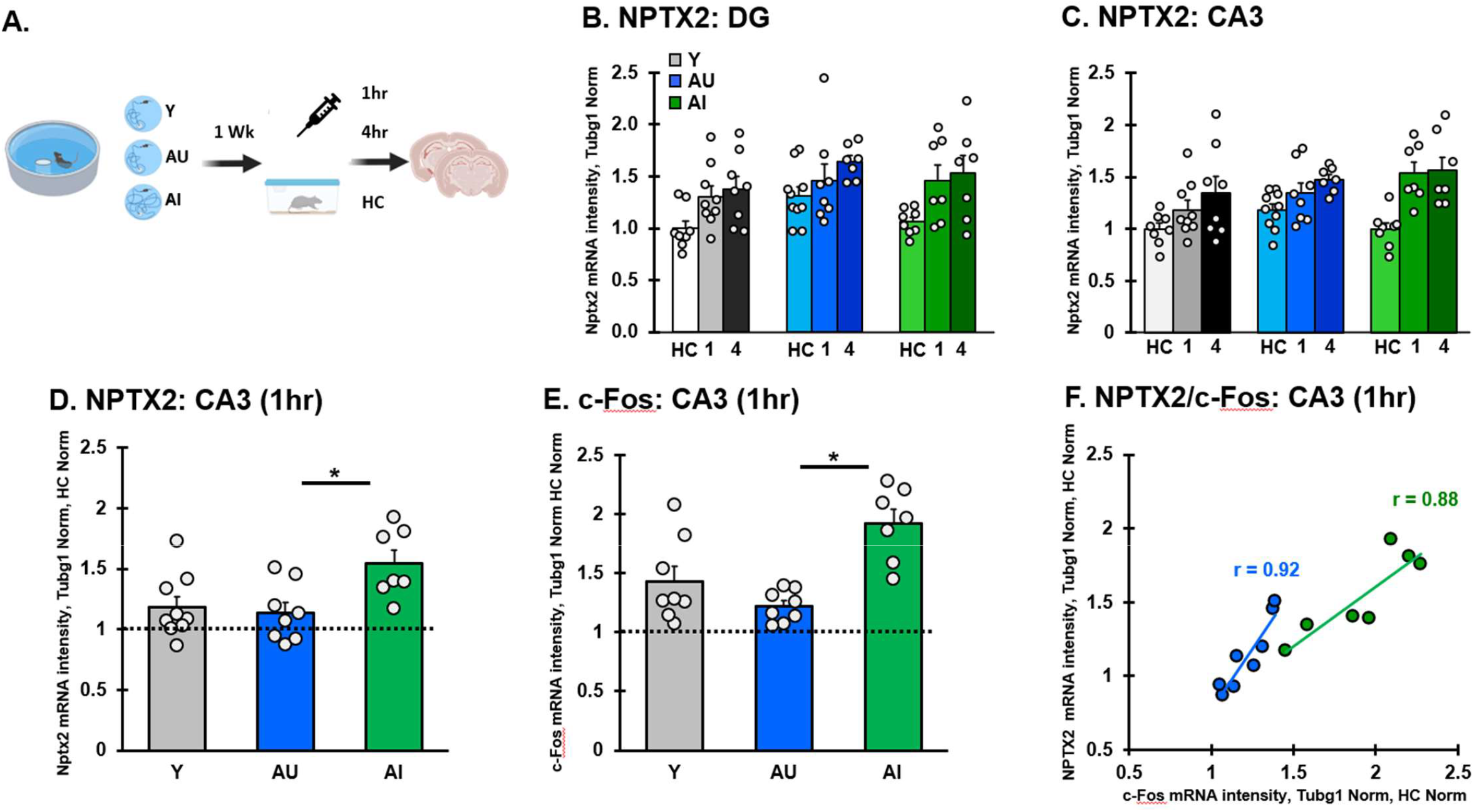
Pharmacological induction of neural activity by pilocarpine administration results in increased NTPX2 transcription in young and aged animals. A) Schematic of pilocarpine gene induction protocol. 1 week following MWM behavioral characterization rats were assigned to treated or untreated (homecage) control groups and brains were collected either 1 hour or 4 hours post pilocarpine treatment to assess short term and long term gene induction following activity. (B, C) Quantification of induced NPTX2 mRNA by in situ hybridization in brain sections of Y, AU, and AI rats in control and activated conditions in DG and CA3. NPTX2 mRNA shows a time dependent elevation following neural activity induction in all groups. In CA3, AI rats had a greater elevation over homecage levels (dashed line) for NPTX2 mRNA (D) and c-Fos mRNA (E) relative to Y and AU rats. (F) 1 hour post pilocarpine, levels of c-Fos and NPTX2 mRNA were positively correlated in AU and AI groups indicating coupling between NPTX2 gene induction and response to neural activity. Significant difference across groups was determined by 2-way ANOVA. Post hoc significance was determined by t-test as indicted on the graphs: ∗p < 0.05; ∗∗p < 0.01. Individual data points represent individual subject values and bar values represent group means ±SEM. Schematic created with BioRender.com.

**Figure 6.**
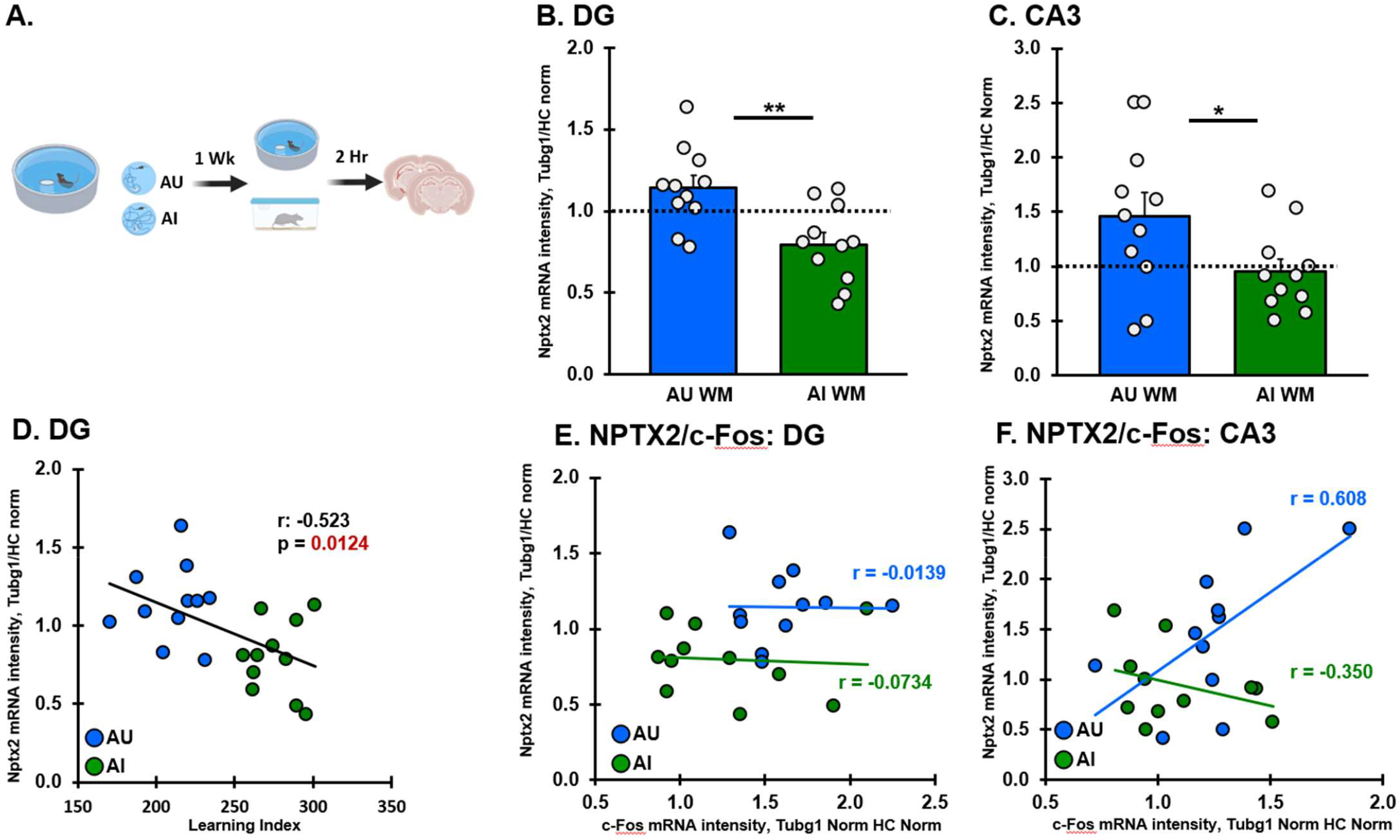
Behavioral induction of neural activity by MWM training results in increased NTPX2 transcription in AU rats relative to AI. A) Schematic of MWM gene induction protocol. 1 week following MWM behavioral characterization rats were assigned to MWM or untreated (homecage) control groups and brains were collected 2 hours post behavior. (B, C) Quantification of induced NPTX2 mRNA by in situ hybridization in AU and AI rats relative to homecage controls in DG and CA3 shows a lack of behavioral induction of NPTX2 in AI rats. Dashed line indicates homecage NPTX2 mRNA intensity level. (D) NPTX2 mRNA behavioral induction was negatively correlated with MWM learning index in the DG subfield. (E) NPTX2 mRNA was not correlated with c-Fos mRNA levels in the DG. (F) In CA3, NPTX2 mRNA levels were positively correlated with c-Fos induction in AU rats and negatively correlated in AI. Significant difference across groups was determined by 1-way ANOVA. Post hoc significance was determined by t test as indicted on the graphs: ∗p < 0.05; ∗∗p < 0.01. Individual data points represent individual subject values and bar values represent group means ±SEM. Schematic created with BioRender.com.

Given the baseline increase in NPTX2 levels in AU animals, it is important to understand the group differences in DG and CA3, and to examine the relative increases in gene expression we normalized NPTX2 expression levels from 1 hour post pilocarpine administration to each group’s average homecage expression level. CA3 was the only subfield to show a significant effect of group (Fig 5D, p=0.014), such that AI exhibited increased activity-induced expression of NPTX2 compared to AU (p=0.007) as well as Y (p=0.013), but expression levels in Y and AU were similar (p=0.709) Thus, both AU and AI were able to respond to an activating stimulus by increasing NPTX2 expression over homecage levels, but while AU rats showed expression increases on par with Y, AI rats showed elevated relative induction of NPTX2 mRNA. In a previous study employing pilocarpine, we identified elevated pilocarpine-induced expression levels in AI rats of the immediate early gene, c-Fos, commonly used as an indicator of neural activity (35). We thus examined c-Fos expression in the current pilocarpine induced cohort, and confirmed higher AI expression levels in CA3 (Fig. 5E, p<0.0001) and DG (p=0.004) further supporting subfield specific enhancement of excitability in the hippocampi of AI rats. To determine if the levels of NPTX2 were related to neuronal activity levels, we examined the relationship between NPTX2 and c-Fos in individual rats in the CA3 subfield (Fig. 5F). NPTX2 and c-Fos expression were highly correlated across all aged rats (r=0.89, p<0.0001), and they were also correlated within animals of the same group (AU: r=0.92, p=0.001; AI: r=0.88, p<0.004). AI rats show higher activity-induced expression of both genes, suggesting the higher neural activity in AIs results in elevated NPTX2. Together, these data demonstrate that the cellular mechanisms needed to support activity-dependent induction of NPTX2 are present in AI rats, but that levels of induction are elevated in AI alongside excessive neuronal activity. For aged unimpaired rats, activity-dependent induction levels were on par with young rats, which does not support the hypothesis that elevated basal NPTX2 levels serve to “jump start” the neuronal induction responses. Rather, AU NPTX2 levels are induced to the same degree as young above the elevated homecage levels. These data are more consistent with the idea that the elevated homecage mRNA expression reflects higher basal levels of homeostatic inhibition, rather than contributing to activity-dependent expression.

### Behaviorally induced expression

To examine behaviorally induced NPTX2 expression, a new set of AU and AI rats were given a second water maze training opportunity that included a new maze with distinct cues and a visible escape platform to equate behavior across both AU and AI. This training occurred with a compressed time schedule such that rats completed 8 training trials within 1 hour and gene expression measured 2 hours from the beginning of training. Previous experiments using this protocol demonstrated activation of learning-related genes and distinct gene expression profiles between AU and AI rats (37). Here, AU rats showed greater behaviorally induced NPTX2 expression relative to homecage controls in DG and CA3, while NPTX2 was reduced below homecage levels in AI rats (Fig. 5B, C). Intriguingly, while the ratio of c-Fos induction relative to NPTX2 was similar in AU and AI in DG (Fig. 5E), there was a striking dissociation between c-Fos and NPTX2 in CA3 (Fig. 5F), such that AU animals recruited NPTX2 in proportion to c-Fos, while AI animals recruited less NPTX2 relative to c-Fos in response to activity. The CA3 subfield, which is known to be involved in pattern completion, has been demonstrated to be hyperactive in aged impaired animals and this hyperactivity is related to a maladaptive bias towards pattern completion. In this context, the potential impact of down regulation of NPTX2 in the face of higher neural engagement implicates a disruption of homeostatic balance that may limit the ability of AI rats to form place fields or remap appropriately in new contexts in downstream CA1 (20).

## Discussion

Distinguishing normal brain aging from the earliest stages of neurodegeneration remains a major challenge in human studies, in part due to age-associated comorbidities that obscure underlying mechanisms. In this study, we leveraged an established rat model of cognitive aging to examine how expression of the activity-dependent gene NPTX2 varies with individual differences in memory function. Aged rats with preserved spatial memory (AU) exhibited a distinct pattern of NPTX2 dynamics compared to both young adults and aged impaired (AI) animals, characterized by elevated baseline expression and appropriate upregulation during learning. In contrast, memory-impaired aged rats showed reduced basal NPTX2 mRNA and protein levels across key hippocampal subfields, and failed to mount a behaviorally driven transcriptional response. These findings support the idea that disrupted NPTX2 regulation marks a specific neurobiological phenotype of cognitive decline in aging, one that may render hippocampal circuits more vulnerable to degeneration, and, conversely, that successful cognitive aging may be supported by adaptive molecular and circuit-level mechanisms that preserve excitatory-inhibitory balance.

One of the most striking findings was elevated NPTX2 mRNA expression in AU rats relative to both AI and young (Y) adults under homecage conditions. In aged rats, higher NPTX2 mRNA levels in DG and CA3 were significantly associated with better cognitive performance, and NPTX2 and GluA4 protein levels in CA1 also covaried both with each other and with memory outcomes. As NPTX2 is trafficked to axon terminals and acts presynaptically to stabilize GluA4 AMPA receptors at PV interneuron synapses (12, 13), these data imply that elevated NPTX2 in AU rats enhances homeostatic inhibitory tone. This aligns with earlier studies showing increased inhibitory control and reduced CA3 hyperactivity in aged rats with preserved memory (18, 26, 29, 38). Our results suggest that elevated NPTX2 in AU rats may reflect an adaptive shift in hippocampal circuit properties that restrains excitability and supports memory function. Prior evidence that AU rats show greater recruitment of inhibitory markers in CA3 during behavior (37, 38) further supports the idea that higher NPTX2 levels contributes to circuit-wide enhancement of inhibitory tone to support the computational functions of the aged hippocampus. Despite lower basal levels, AI rats retained the capacity to upregulate NPTX2 following pharmacological neuronal activation. As an immediate early gene, NPTX2 mRNA is rapidly produced in response neuronal activity and participates in the generation of synapse remodeling and homeostatic plasticity in synaptic circuits throughout the brain (11, 14). Notably, aging did not impair this response; both AU and AI animals exhibited strong NPTX2 mRNA induction following pilocarpine administration, indicating that the gene’s basic molecular responsiveness remains intact with age, consistent with electrical induction of LTP in Fischer rats (39). Interestingly, with this experimental procedure, AI rats showed even greater activity-induced NPTX2 expression in CA3 than AU or young animals. The tight correlation in all animal groups between NPTX2 and c-Fos expression, a measure of neuronal activity, suggests that greater NPTX2 induction occurs with greater neural activity, mirroring prior reports of elevated c-Fos and hyperexcitability in AI rats (35). This disproportionate induction relative to baseline suggests that AI hippocampal networks are prone to excessive excitation, likely reflecting the homeostatic imbalance in inhibitory tone. In this context, NPTX2 mRNA may reflect compensatory over-engagement rather than effective inhibition. This contrasts with AU rats, where elevated baseline levels of NPTX2 likely promote sustained inhibitory tone and enable more efficient responses to behavioral demands.

Most importantly, the behavioral-induction experiment provides the clearest evidence that AI and AU animals diverge in how they engage inhibitory control during learning. After a rapid water-maze training protocol in a novel environment, AU rats upregulated NPTX2 in both DG and CA3, whereas AI rats showed a decrease in NPTX2 despite robust c-Fos induction. This decoupling between activity and NPTX2 transcription was most pronounced in CA3, a region whose hyperactivity has been implicated in maladaptive pattern completion during aging (20, 40). These findings suggest that physiologically relevant hippocampal activation is insufficient to induce NPTX2 in AI rats, despite their capacity to express NPTX2 in response to broad, non-specific neural activation. This failure to respond appropriately to behaviorally relevant stimuli likely reflects the underlying state of the network, which influences the coupling between neuronal activity and transcriptional responses (41). Enhanced inhibition in AU rats, as indicated by elevated basal NPTX2 levels in the current study and increased expression of inhibition-related genes in CA3 following behavior (37), may support a more balanced and dynamic network. In contrast, impaired signal sensitivity in AI animals may reflect, at least in part, the circuit hyperactivity observed in both rodent and human studies (30, 31). Supporting this interpretation, recent work using hippocampal slice recordings has shown that diminished recruitment of PV interneurons in response to perforant path stimulation in AI rats can be rescued by NPTX2 expression in the lateral entorhinal cortex, an intervention sufficient to improve cognitive performance. Together, these results support a model in which enhanced inhibition in aging, both at baseline and in response to behavioral demands, is essential for maintaining hippocampal network homeostasis and enabling the recruitment of appropriate memory mechanisms in response to sensory input.

Furthermore, western blot analyses of homecage animals showed elevated NPTX2 protein in CA1, but not CA3, despite increased NPTX2 mRNA in CA3 of AU rats. This suggests that NPTX2 is synthesized in CA3 and trafficked to axon terminals in CA1, where it may modulate postsynaptic targets. In CA1 NPTX2 levels strongly correlated with GluA4, a calcium-permeable AMPA subunit enriched at PV interneuron synapses. This relationship supports the idea that NPTX2 enhances GluA4 stabilization on PV interneurons, promoting their activity (12, 13). Thus, failure of AI rats to induce NPTX2 in CA3 during learning may reduce delivery of NPTX2 to CA1, weakening PV interneuron recruitment and disrupting inhibitory control over CA1 pyramidal neurons. This impairment in feedforward inhibition may reduce the fidelity of CA1 output, limiting the hippocampus’s capacity to flexibly encode new spatial representations and contributing to memory impairments in aging.

In aged Long Evans rats, the functional and molecular condition of the MTL memory circuit shares key features with individuals diagnosed with the earliest phases of cognitive impairment and early amnestic MCI, a symptomatic phase of AD that emerges prior to widespread neurodegeneration (27, 42). Impaired spatial navigation and episodic memory, neuronal hyperactivity, and disrupted circuit connectivity are observed in both individuals with aMCI and aged rats with cognitive impairment relative to mature young adults. Thus, this aged outbred rodent strain captures physiological features of MTL functioning associated with age-related cognitive decline and prodromal AD. Building on evidence linking hippocampal hyperactivity to cognitive impairment in aging (18, 20, 30, 31, 35) and enhanced inhibitory recruitment to aged individuals with preserved cognitive function (29, 38), we found that NPTX2 expression tracks the individual differences in cognitive status, particularly in CA3 and DG, regions vulnerable to age-related shifts in E/I balance. Importantly, AU rats exhibited elevated NPTX2 expression under basal conditions and appropriate activity-dependent regulation during behavioral learning, while AI rats displayed reduced basal levels and aberrant expression patterns in response to behavioral activation. Notably, these findings, in particular, suggest NPTX2 as a key molecular correlate of preserved inhibitory tone and a potential contributor to resilience against age-related memory decline.

Together, our findings provide mechanistic insight into the clinical observation that reduced NPTX2 levels in aging and AD are associated with cognitive decline observed in those conditions. Human studies consistently link lower NPTX2 in CSF and brain tissue with impaired cognition and reduced hippocampal volume, and longitudinal decreases in NPTX2 predict progression to MCI even in the absence of classic AD pathology (6, 8, 10, 43). Longitudinal data further reveal that declines in CSF NPTX2 precede overt neurodegeneration and may predict cognitive decline independently of amyloid or tau pathology (8, 10). However, these studies cannot distinguish whether NPTX2 loss is a driver or consequence of neural dysfunction. Using a reverse translational approach in this rat model of healthy cognitive aging, we show that low NPTX2 expression in aged, cognitively impaired animals occurs alongside disrupted activity-dependent regulation and a failure to sustain inhibitory tone in hippocampal circuits. In contrast, preserved memory in aged unimpaired rats is associated with elevated basal NPTX2 levels and appropriate upregulation during behavioral learning, suggesting that NPTX2 actively contributes to maintaining excitatory-inhibitory balance. Our results indicate that NPTX2 is more than a passive marker of synaptic health, and may be a functional mediator of inhibitory control and cognitive resilience *in vivo*. This model clarifies the biological significance of NPTX2 reductions observed in clinical studies and supports its use as a prognostic indicator and potentially an early therapeutic target for age-related cognitive decline.

## Materials and Methods

### Subjects

Aged LE rats were obtained at 8-9 months of age from Charles River Laboratories (Raleigh, NC) and housed in a vivarium at Johns Hopkins University until 24-26 months of age. Young rats obtained from the same source were tested around 4-6 months of age. All rats were individually housed at 25°C and maintained on a 12 hr light/dark cycle. Food and water were provided ad libitum unless indicated otherwise. Aged rats were examined for health and pathogen-free status throughout the study, as well as necropsies at the time of sacrifice. Rats that showed impaired health or disabilities that could impact behavioral performance (e.g. poor eyesight, clinical evidence of renal impairment, pituitary or other tumors) were excluded from the analyses. All procedures were approved by the Johns Hopkins University Institutional Animal Care and Use Committee in accordance with the National Institutes of Health directive.

### Behavioral Characterization

Young adult (8–9 months) and aged rats (24 months) were tested in an assessment of hippocampal function prior to any subsequent manipulations. The background behavioral assessment used a well-established Morris water maze protocol as described in detail elsewhere (21, 44). This protocol was designed to examine spatial reference memory with sparse training (3 trials per day) at 24-h intervals. Rats were trained for 8 days to locate a camouflaged escape platform that remained at the same location throughout training in a water maze surrounded by curtains with fixed cues. Every sixth trial consisted of a probe trial (no escape platform for the first 30s of the trial) that served to assess the development of a spatially localized search. Learning Index (LI) scores were derived from each rat’s proximity to the platform during the four probe trials. The proximity measure was obtained by sampling the position of the animal in the maze (10 times per second) to provide a record of its distance from the escape platform in 1-s averages. The learning index is the sum of weighted proximity scores obtained during probe trials, with low scores reflecting a more accurate search and indicating better retention of the platform location. A learning index cutoff was used to identify aged rats as Aged Unimpaired (AU, LI<240) or Aged Impaired (AI, LI>240) with higher scores representing worse performance and reflecting scores that fall outside the normative range collected from young adult LE rats over many years. Visible platform (cue) training was used to assess the sensorimotor and motivational status of the rats. Only rats with successful cue training performance were included in the present study. After behavioral characterization and cue training, animals were selected for inclusion in further studies to create balanced groups of young, aged impaired, and aged unimpaired subjects in five tissue libraries (Homecage, Pilocarpine, Water Maze, RNAseq, Western). LI distribution for each tissue library is shown in Supplemental Fig. 1. Subjects used to assess homecage (basal) gene expression through RNA-seq or *in situ* hybridization remained in their home cages for at least 10 days after behavior assessment prior to sacrifice. After excluding subjects with pituitary (or other brain) tumors, adverse reactions to pilocarpine (death, n=5) or outlier expression values for control genes (n=3, 4 hr pilocarpine treated subjects, n=1 water maze subject, n=1 DESeq subject), data from 201 rats were included across all studies.

### Basal gene expression (Homecage tissue library)

For basal *in situ* hybridization, 16 Y, 19 AU, and 19 AI rats were taken directly from the homecage prior to sacrifice (see Supp. Fig. 1 for LI distribution). Rats were anesthetized with isoflurane and perfused transcardially with ice cold 0.1-M phosphate buffered saline, followed by 4% paraformaldehyde in 0.1-M phosphate buffer. Brains were harvested and post-fixed in paraformaldehyde overnight and then in 16% sucrose with 4% paraformaldehyde for another 48 hours. Homecage controls for the pilocarpine tissue library (described below) were included in the homecage tissue library for assessing basal NPTX2 expression (Fig. 3).

### Pharmacological induction of neural activity (Pilocarpine tissue library)

25 Y, 26 AU, and 23 AI animals were selected and assigned to homecage (control), 1-hour post induction, and 4-hour post induction groups (see Supp. Fig. 1 for LI distribution). As in (36) and (35), a low, subconvulsive dose of pilocarpine was administered systemically to induce widespread neural activity across the brain. Pilocarpine, a muscarinic acetylcholine receptor agonist, modulates both inhibitory and excitatory synaptic transmission with a net excitatory effect on brain activity that reflects the condition of the activated networks (45, 46). At the time of pharmacological treatment, rats were injected with pilocarpine (25 mg/kg in a volume of 1 mL/kg in saline) and left undisturbed in their home cage for either 1 hour or 4 hours to assess short term and delayed gene induction. Rats were monitored for peripheral cholinergic symptoms (tremor and lacrimation) by a trained observer. No overt seizures were observed between pilocarpine administration and sacrifice. Aged and young rats exhibited similar symptoms over similar durations. One hour or 4 hours after pilocarpine injection, the rats were anesthetized with isoflurane and perfused as indicated above.

### Behavioral gene expression induction protocol (Water Maze tissue library)

22 AU and 23 AI animals were selected and assigned to homecage (control) or 1-hour post behavioral treatment groups (see Supp. Fig. 1 for LI distribution). 4-6 weeks after behavioral characterization, all rats were given a single training session (8 trials with 8-min ITI) in a new water maze environment located at a different site as described in detail in Haberman et al 2008. The maze environment contained prominent extramaze spatial cues and a visible escape platform that remained at the same location for all training trials. Intact escape performance was used to ensure the absence of sensorimotor and motivational deficits. One hour after the last training trial, all rats were given 90-sec probe trial without the escape platform. Data were analyzed with a video tracking system (HVS Image Analyzing VP-116) and an IBM PC computer with software developed by HVS Imaging (Hampton, UK). Directly after probe trial completion, rats were anesthetized with isoflurane and perfused as indicated above.

### RNA sequencing (RNAseq tissue library)

8 Y, 8 AU, and 8 AI subjects were. RNA sequencing was performed on RNA extracted from CA3 tissue dissected from 800µM transverse slices throughout the entire rostral-caudal extent of the hippocampus. All RNA samples were subjected to quality control analysis via Agilent 2100 bioanalyzer and samples with the best quality (24 samples in total with RNA integrity numbers above 8.5) were submitted for sequencing through Center for Computational Genomics and Next Generation Sequencing Center within the Sidney Kimmel Comprehensive Cancer Center at Johns Hopkins University. mRNA was prepared for sequencing using Illumina TruSeq Strand Specific Total RNA sample prep and sequencing was performed on Illumina HiSeq2500 high output mode with paired end reads. Alignments were performed using RSEM to the Rn5 rat genome and differential expression determined with DEseq. Total reads ranged from 62-86 million per sample while aligned reads ranged from 21 – 34 million per sample (approximately 40% of total reads). There were no differences between age or cognitive groups in total or aligned reads. Graphed gene expression data is derived from DEseq generated expected gene counts normalized to total reads and then normalized to the average of Y rats for each gene. Statistical differences are reported as raw p-values and a q-value adjusted for multiple comparisons. Q<0.1 was considered significant.

### MTL subfield dissections (Western tissue library)

Two weeks after completion of behavioral testing, rats (5 Y, 12 AU, and 13 AI (5 female)) were deeply anesthetized with isoflurane and sacrificed by rapid decapitation (see Supp. Fig. 1 for LI distribution). Brains were immediately extracted and placed in ice-cold phosphate-buffered saline (PBS). The hippocampus was separated from the brain and CA1, CA3, and DG were microdissected by hand from 800µM transverse sections of the hippocampus along its entire longitudinal extent as previously described (47). Rat brain tissue were lysed with RIPA buffer previously described (48). Protein extracts were separated by 4 to 12% SDS-PAGE (Thermo Fisher Scientific; WG1403BOX), and transferred to polyvinylidene difluoride membranes (Millipore, IPFL00010). Membranes were blocked with 5% nonfat milk in tris-buffered saline with 0.1% Tween 20 (TBST) for 1 hour at room temperature. Primary antibodies in TBST were incubated overnight at 4°C. After washes with TBST to remove primary antibodies, membranes were incubated with HRP-conjugated secondary antibodies for 1 hour at room temperature. Additional washes with TBST to remove secondary antibodies were done, then membranes were exposed with West Pico ECL, and GluA4 was additionally exposed with West Femto ECL. Immunoreactivity was detected using ChemiDoc system (Bio-Rad) and analyzed with Image Lab software (Bio-Rad). PSD95 and actin were used as loading controls. Rabbit anti-NPTX2 antibody (1:1,000) was previously described (14). Commercial antibodies included Rabbit anti-GluA4 (1:1,000) (Millipore Sigma, Cat. No. AB1508). Mouse anti-PSD95 (1:1,000) (Thermo Fisher Scientific, Cat No. MA1-046; RRID: AB_2092361); Mouse anti-actin (1:10,000) (Sigma-Aldrich, Cat.No. A2228; RRID: AB_476697). Secondary antibodies (1:10,000) were ECLTM (enhanced chemiluminescence) anti-mouse immunoglobulin G (IgG) horseradish peroxidase (HRP) and ECLTM anti-rabbit IgG HRP (GE Healthcare, Cat. No. NA931V and NA934V, respectively). Western Blot detection was performed using SuperSignal West Pico (Luminol Enhancer Solution, Cat. No. 1863098; Stable Peroxide Solution Cat. No. 1863099) and SuperSignal West Femto (Luminol Enhancer Solution, Cat. No. 1859022; Stable Peroxide Solution, Cat. No. 1859023) from Thermo Fisher Scientific.

### In situ hybridization

NPTX2 and c-Fos mRNA expression was measured by quantitative *in situ* hybridization with Tubg1 expression used to control for systematic differences in tissue quality. Brains were cut coronally at 40 um using a freezing microtome in a 1 in 20 series across the entire extent of the medial temporal lobe. Sections were stored in paraformaldehyde at 4ºC until processed for *in situ* hybridization. Probe generation and *in situ* hybridization were performed as in Haberman et al. (2011). Briefly, brain sections matched for number and location were hybridized overnight at 60ºC in buffer containing a 35S-UTP-labeled riboprobe generated using the MAXIscript kit (Ambion). *In situ* probe sequences were either PCR amplified from whole hippocampal cDNA with primers incorporating T7 and SP6 RNA polymerase binding sites or PCR-derived amplicons were cloned in pGEM plasmids and digested with appropriate restriction enzymes. Probe sequences were as follows: Nptx2, nts 1551 to 1903 of Genbank seq S82649.1; Tubg1, nts1252-1601 of GenBank seq NM_145778.2; and c-Fos, nts 670 to 1043 from GenBank seq NM_022197.2. Each probe was shown to have little to no homology to any other rat gene based on a BLAST search of each fragment sequence. Hybridized sections were then extensively washed and mounted onto slides. Mounted, dried sections were then exposed in a phosphorimager cassette for several days and scanned using Typhoon 5 Variable Mode Imager (GE Healthcare, PA, USA). Brain regions of interest were outlined by hand and matched for level along the anterior-posterior axis and quantified, blind to experimental conditions, using ImageQuant (GE Healthcare, PA, USA). The areas analyzed included dentate gyrus (DG), CA1, CA2 and CA3 subfields of the hippocampus (extending from approximately 2.3 mm to 4.16 mm relative to bregma). Radioactive standards exposed at the same time as the brain sections ensured that section intensity was within the linear range. The intensity level of mRNA labeling of at least 4 sections for each area of interest was averaged to obtain a single score for each rat. In rare cases, tissue integrity was compromised such that quality data could not be obtained, resulting in exclusion of an animal for that brain region. This is reflected in the N values for each analysis. To maintain quality, *in situ* hybridizations were performed in batches with separate scans. Average intensities from each scan were normalized to standards, and z-scored before combining into a single dataset. Z-scored values were back transformed and normalized to TubulinG1 (Tubg1) mRNA intensities in dorsal hippocampus (Supp. Figure 2). Basal expression data included subjects from an independent homecage set as well as the homecage control subjects from the pilocarpine induction study. Significance was determined by univariate 2 factor analysis, analysis of variance followed by Tukey’s post hoc for independent samples and T-test as appropriate. All analyses were performed with IBM SPSS statistics software.

### Constructing the Protein–Protein Interaction (PPI) Networks

The search tool for retrieval of interacting genes (STRING) (https://string-db.org) database, which integrates both known and predicted PPIs, can be applied to predict functional interactions of proteins (49). To seek potential interactions between differentially expressed genes in cognitively characterized aged rats, the STRING tool was employed with the search term ‘NPTX2’. Active interaction sources, including text mining, experiments, databases, and co-expression as well as species limited to “Homo sapiens” and an interaction score > 0.4 were applied to construct the PPI networks. In the network, the nodes correspond to the proteins and the edges represent the interactions. The PPI network was used for identification of potential NPTX2 partner proteins and not for quantitative analysis.

## Acknowledgments

The authors would like to acknowledge Rob McMahan and Jala Atufa for assistance with behavioral experiments and Arnold Bakker for consultation on the manuscript. This work was supported by National Institute on Aging grant P01AG009973.

