## Supplemental Figures for "Altered NPTX2 dynamics associated with impaired cognitive aging"

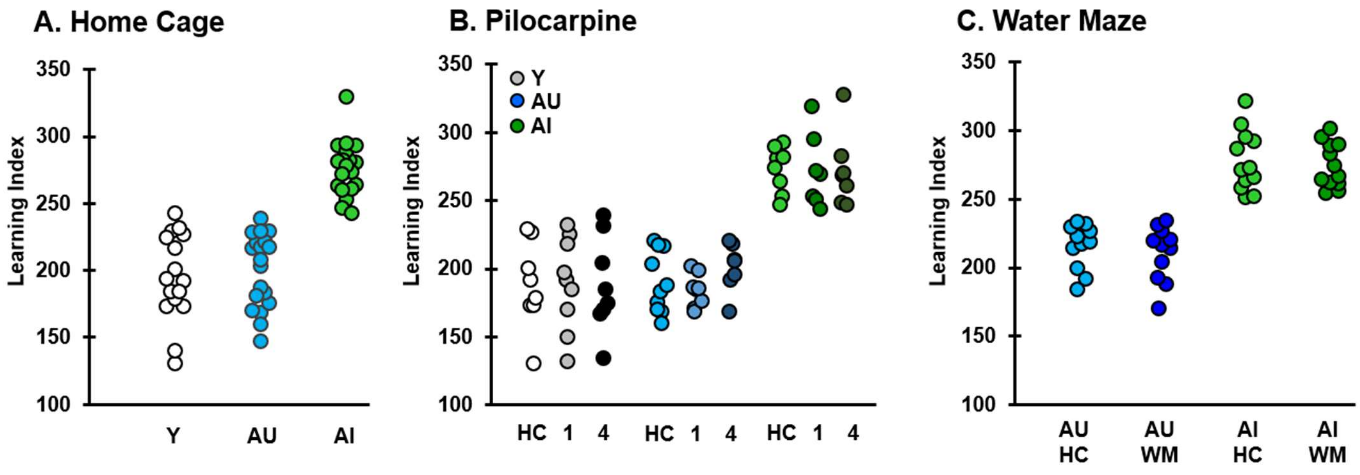


**Supplemental Figure 1.** MWM learning index values for all subjects in experimental tissue libraries.

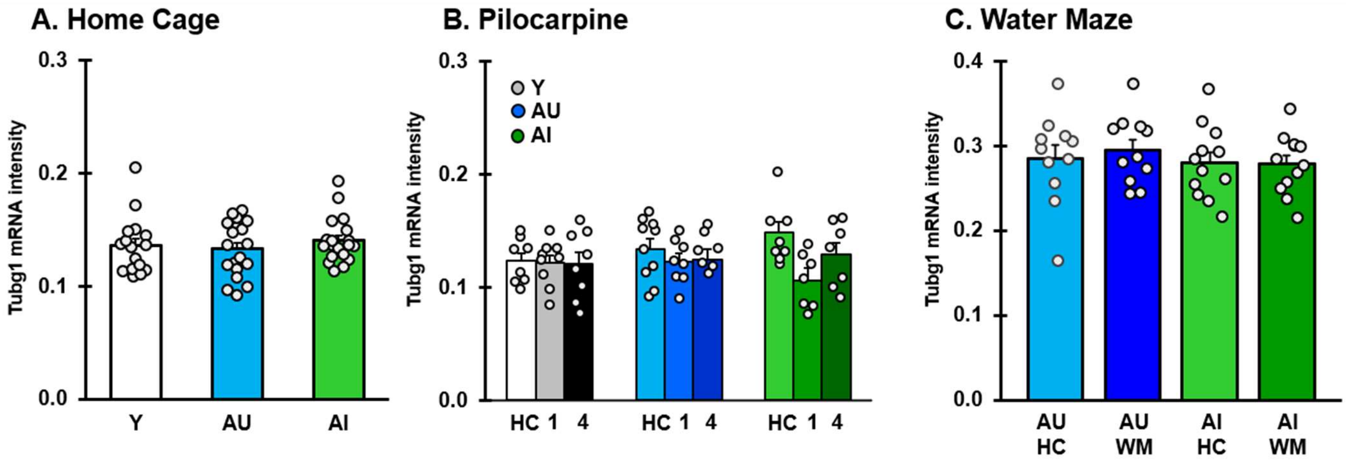


**Supplemental Figure 2**. TubulinG1 (Tubg1) mRNA control gene expression for all subjects.

| | | | | Group statistics | | | One-way ANOVA | | | | | | Tukey HSD (pLL, UL) | | |
|---|---|---|---|---|---|---|---|---|---|---|---|---|---|---|---|
| Experiment | Measure | Region | Group | n | Mean | SEM | SS | df | MS | F | Sig. | η2 | Y v AU | Y v AI | AU v AI |
| Homecage | NPTX2 | DG | Y | 16 | 1.0904 | 0.0567 | 0.907 | 2 | 0.4535 | F(2,53)=12.314 | 4.33E-05 | 0.325651 | 0.000(-0.470,-0.156) | 0.270(-0.259,0.055) | 0.004(0.061,0.361) |
| | | | AU | 19 | 1.4033 | 0.0426 | | | | | | | | | |
| | | | AI | 19 | 1.1923 | 0.0377 | | | | | | | | | |
| | | CA3 | Y | 16 | 0.9454 | 0.0325 | 0.434 | 2 | 0.217 | F(2,53)=10.880 | 0.000116 | 0.299068 | 0.001(-0.306,-0.074) | 0.996(-0.119,0.112) | 0.00(0.0753,0.297) |
| | | | AU | 19 | 1.1352 | 0.0360 | | | | | | | | | |
| | | | AI | 19 | 0.9493 | 0.0306 | | | | | | | | | |
| | | CA2 | Y | 16 | 1.4471 | 0.0980 | 1.003 | 2 | 0.5013 | F(2,53)=4.562 | 0.015049 | 0.15174 | 0.025(-0.575,-0.0318) | 0.944(-0.308,0.235) | 0.042(0.008,0.527) |
| | | | AU | 19 | 1.7505 | 0.0710 | | | | | | | | | |
| | | | AI | 19 | 1.4833 | 0.0679 | | | | | | | | | |
| | | CA1 | Y | 16 | 0.6893 | 0.0364 | 0.013 | 2 | 0.0066 | F(2,53)=0.316 | 0.730467 | 0.012241 | | | |
| | | | AU | 19 | 0.7027 | 0.0279 | | | | | | | | | |
| | | | AI | 19 | 0.6658 | 0.0377 | | | | | | | | | |
| | | LEC | Y | 15 | 1.0637 | 0.0704 | 0.099 | 2 | 0.0496 | F(2,52)=0.852 | 0.432812 | 0.032943 | | | |
| | | | AU | 19 | 0.9596 | 0.0613 | | | | | | | | | |
| | | | AI | 19 | 0.9792 | 0.0415 | | | | | | | | | |
| | | MEC | Y | 16 | 0.7137 | 0.0437 | 0.001 | 2 | 0.0003 | F(2,53)=0.013 | 0.987416 | 0.000497 | | | |
| | | | AU | 19 | 0.7062 | 0.0415 | | | | | | | | | |
| | | | AI | 19 | 0.7132 | 0.0265 | | | | | | | | | |

| | | | | | Group statistics | | | | Two-way ANOVA (Group, time post treatment) | | | | | | | Tukey HSD | |
|---|---|---|---|---|---|---|---|---|---|---|---|---|---|---|---|---|---|
| Experiment | Measure | Region | Treatment | Group | n | Mean | SEM | Factor | SS | df | MS | F | Sig. | Partial $\eta^2$ | | | |
| Pilocarpine | NPTX2 | DG | HC | Y | 8 | 1.106 | 0.081 | Overall | 3.237 | 8 | 0.405 | F(8,63) = 3.292 | 0.003 | 0.295 | | | |
| | | | | AU | 10 | 1.449 | 0.094 | Group | 0.883 | 2 | 0.442 | F(2,63) = 3.593 | 0.033 | 0.102 | Y vs AU: P=0.045 | Y vs AI: P=0.492 | AU vs AI: P=0.443 |
| | | | | AI | 8 | 1.178 | 0.048 | Treatment | 2.409 | 2 | 1.204 | F(2,63) = 9.802 | 0.000 | 0.237 | 0 vs 1Hr: P=0.013 | 0 vs 4Hr: P=0.000 | 1 vs 4Hr: P=0.483 |
| | | | 1Hr | Y | 9 | 1.442 | 0.110 | Interaction | 0.157 | 4 | 0.039 | F(4,63) = 0.320 | 0.863 | 0.020 | | | |
| | | | | AU | 8 | 1.615 | 0.175 | | | | | | | | | | |
| | | | | AI | 7 | 1.611 | 0.168 | | | | | | | | | | |
| | | | 4Hr | Y | 8 | 1.526 | 0.135 | | | | | | | | | | |
| | | | | AU | 7 | 1.812 | 0.068 | | | | | | | | | | |
| | | | | AI | 7 | 1.689 | 0.187 | | | | | | | | | | |
| | | CA3 | HC | Y | 8 | 0.934 | 0.051 | Overall | 2.45 | 8 | 0.306 | F(8,63) = 5.022 | 0.000 | 0.389 | | | |
| | | | | AU | 10 | 1.103 | 0.053 | Group | 0.421 | 2 | 0.211 | F(2,63) = 3.454 | 0.038 | 0.099 | Y vs AU: P=0.167 | Y vs AI: P=0.078 | AU vs AI: P=0.903 |
| | | | | AI | 8 | 0.930 | 0.056 | Treatment | 1.847 | 2 | 0.923 | F(2,63) = 15.141 | 0.000 | 0.325 | 0Hr vs 1 Hr: P=0.002 | 0Hr vs 4Hr: P=0.000 | 1Hr vs 4Hr: P=0.293 |
| | | | 1Hr | Y | 9 | 1.106 | 0.082 | Interaction | 0.329 | 4 | 0.082 | F(4,63) = 1.347 | 0.000 | 0.079 | | | |
| | | | | AU | 8 | 1.253 | 0.094 | | | | | | | | | | |
| | | | | AI | 7 | 1.435 | 0.100 | | | | | | | | | | |
| | | | 4Hr | Y | 8 | 1.259 | 0.144 | | | | | | | | | | |
| | | | | AU | 7 | 1.379 | 0.043 | | | | | | | | | | |
| | | | | AI | 7 | 1.459 | 0.122 | | | | | | | | | | |
| | | | | | | | | | | | | One-way ANOVA | | | | Tukey HSD (p(CI LL, CI UL)) | |
| | | | | | | | | | SS | df | MS | F | Sig. | Partial $\eta^2$ | Y vs AU | Y vs AI | AU vs AI |
| | cFOS | CA3 | 1Hr | Y | 8 | 1.957 | 0.172 | | 1.901 | 2 | 0.951 | F(2,22)=11.988 | 0.000 | 0.545 | 0.312(-0.145, 0.568) | 0.008(-0.858, -0.120) | 0.000((-1.069, -0.332) |
| | | | | AU | 8 | 1.842 | 0.071 | | | | | | | | | | |
| | | | | AI | 7 | 2.387 | 0.145 | | | | | | | | | | |

| | | | | | | Group statistics | | Independent Sample Ttest | | | Cohen's d (CI UL, CI LL) |
|---|---|---|---|---|---|---|---|---|---|---|---|
| Experiment | Measure | Region | Treatment | Group | n | Mean | SEM | df | t | Sig. | |
| Water maze | NPTX2 | DG | WM | AU | 11 | 1.143 | 0.0731 | 20 | 3.413 | 0.003 | 1.455(0.493,2.390) |
| | | | | AI | 11 | 0.796 | 0.0708 | | | | |
| | | CA3 | WM | AU | 11 | 1.464 | 0.2106 | 20 | 2.152 | 0.044 | 0.918(0.025,1.790) |
| | | | | AI | 11 | 0.949 | 0.1134 | | | | |

**Supplemental Figure 3.** Full statistics for homecage, pilocarpine gene induction, and water maze gene induction tissue sets.
